# Enzyme catalysis prior to aromatic residues: reverse engineering of a dephosphoCoA kinase

**DOI:** 10.1101/2020.11.11.377994

**Authors:** Mikhail Makarov, Jingwei Meng, Vyacheslav Tretyachenko, Pavel Srb, Anna Březinová, Valerio Guido Giacobelli, Lucie Bednárová, Jiří Vondrášek, A. Keith Dunker, Klára Hlouchová

**Affiliations:** Department of Cell Biology, Faculty of Science, Charles University, Biocev, Prague 2, Czech Republic; Department of Biochemistry, Faculty of Science, Charles University, Hlavova 8, 128 00, Prague 2, Czech Republic; Center for Computational Biology and Bioinformatics, Indiana University School of Medicine, Indianapolis, Indiana, USA; Institute of Organic Chemistry and Biochemistry IOCB Research Centre & Gilead Sciences, Academy of Sciences of the Czech Republic, Flemingovo nám. 2, 166 10, Prague, Czech Republic; Proteomics Core Facility, BIOCEV, Faculty of Science Charles University, Prague, Czech Republic

**Author notes:** shared authorship (Mikhail Makarov, Jingwei Meng).

**Keywords:** genetic code evolution, protein disorder, catalysis evolution, aromatic amino acids

## Abstract

It is well-known that the large diversity of protein functions and structures is derived from the broad spectrum of physicochemical properties of the 20 canonical amino acids. According to the generally accepted hypothesis, protein evolution was continuously associated with enrichment of this alphabet, increasing stability, specificity and spectrum of catalytic functions. Aromatic amino acids are considered the latest addition to genetic code.

The main objective of this study was to test whether enzymatic catalysis can spare the aromatic amino acids (aromatics) by determining the effect of amino acid alphabet reduction on structure and function of dephospho-CoA kinase (DPCK). We designed two mutant variants of a putative DPCK from *Aquifex aeolicus* by substituting (i) Tyr, Phe and Trp or (ii) all aromatics (including His), i.e. ∼10% of the total sequence. Their structural characterization indicates that removal of aromatic amino acids may support rich secondary structure content although inevitably impairs a firm globular arrangement. Both variants still possess ATPase activity, although with 150-300 times lower efficiency in comparison with the wild-type phosphotransferase activity. The transfer of the phosphate group to the dephospho-CoA substrate is however heavily uncoupled and only one of the variants is still able to perform the reaction.

Here we provide support to the hypothesis that proteins in the early stages of life could support at least some enzymatic activities, despite lower efficiencies resulting from the lack of a firm hydrophobic core. Based on the presented data we hypothesize that further protein scaffolding role may be provided by ligands upon binding.

**Significance:** All extant proteins rely on the standard coded amino acid alphabet. However, early proteins lacked some of these amino acids that were incorporated into the genetic code only after the evolution of their respective metabolic pathways, aromatic amino acids being among the last additions. This is intriguing because of their crucial role in hydrophobic core packing, indispensable for enzyme catalysis.

We designed two aromatics-less variants of a highly conserved enzyme from the CoA synthesis pathway, capable of enzyme catalysis and showing significant ordering upon substrate binding. To our knowledge, this is the first example of enzyme catalysis in complete absence of aromatic amino acids and presents a possible mechanism of how aromatics-less enzymes could potentially support an early biosphere.

## Introduction

The extant alphabet of canonical amino acids was apparently selected in the first 10-15% of Earth history from a plethora of amino acids (i) available on primordial Earth and (ii) synthesized through gradually developing metabolic pathways (1). Recent analyses reveal that, compared to alternatives, it composes an unusually good repertoire of physical properties (2-4). Even entirely random sequences built from the canonical alphabet give rise to secondary structure-rich proteins (5). Nevertheless, soluble and well-expressing proteins have been successfully recovered from random libraries of simpler alphabet of evolutionary early amino acids (6, 7). However, the stage of the amino acid alphabet evolution at which proteins could have gained dominance in binding and catalysis remains unclear.

Aromatic amino acids are considered among the last additions to the genetic coding system, i.e. to the canonical amino acid alphabet (8, 9). Because of their relatively high redox reactivity, their fixation in the genetic code could be driven by the biospheric oxygen (10). There is recent evidence that for some of the aromatics (Tyr and Trp) this possibly happened even in the post-LUCA period (10, 11, 12). These proposals suggest that there was a time when living cells existed without aromatic amino acids.

Even though different reduced sets (of 7-13) of the amino acid alphabet have been shown or predicted to be sufficient for protein folding and catalysis, to our knowledge, none of the experimental studies recovered enzyme activity in complete absence of aromatics (13-18). Computational inquiry indicates that the aromatics are the strongest structure promoters among the 20 amino acid alphabet (19). This conclusion is consistent with observation that aromatics are mostly clustered within the hydrophobic cores of structured proteins and with quantum chemistry calculations showing the interactions between aromatics stronger and more specific than aliphatic side chains interactions (20). A comparison of the structure/disorder propensities of the 20 amino acids with the chronology of amino acid inclusion into the genetic code indicates that the earliest amino acids are strongly disorder-promoting while the last to be added, e.g. the aromatics, are among the most strongly structure-promoting (9, 19, 21). Indeed, aromatics are heavily under-represented in intrinsically disordered proteins and regions (IDPs and IDRs), i.e. proteins that lack stable 3D structure and yet frequently carry out crucial biological functions, associated with signaling and regulation in particular (22, 23). While some functions can thus be delivered even in lack of tertiary structure, it remains unclear if and how early enzymes could achieve specific catalysis without a stable hydrophobic core and aid of aromatic residues.

Here, we perform an analysis of structure/function consequences of amino acid reduction by aromatic amino acids. As an exemplary target, we choose a highly conserved metabolic enzyme from a hyperthermophilic bacteria (and hence of potential relevance to early life) - an enzyme that catalyzes the final step of coenzyme A biosynthesis, which is known to be essential for all life and considered among the most ancient cofactors. We present evidence that enzyme catalysis can occur in the absence of aromatic amino acids and a firm hydrophobic core, formation of which can be induced upon ligand binding.

## Results

### Target selection by analysis of LUCA proteins

In order to identify conserved structured protein families, we applied our VSL2B disorder predictor (24) to a collection of last universal common ancestor (LUCA) assigned proteins identified by their ubiquity across all kingdoms of cellular life (25). The non-enzymes, mostly ribosomal or other RNA binding proteins, were all predicted to be massively disordered while the enzymes were predicted to be structured. These modern-day versions of the ancient enzymes all contain multiple aromatic residues, we have been unable to identify a single efficient modern enzyme that lacks multiple aromatic residues. Among the LUCA assigned enzymes identified by Brooks and Fresco, we selected dephospho-CoA kinase (DPCK) for further study because it has the lowest number of aromatic amino acids (25). Other advantages of this choice are that there are multiple 3D structures of different DPCK family members and that the DPCK proteins have relatively small sizes.

### Sequence design, expression and purification of DPCK variants

To evaluate the significance of aromatic amino acids for the structure and function of DPCK, the PDB database was first searched for solved structures of confirmed and putative DPCKs from different thermophilic bacterial species (Supplementary Table S1). An initial test of expression, solubility, ease of large-scale purification and DPCK activity lead to selection of a putative DPCK from *Aquifex aeolicus* (PDB ID: 2IF2) for this experiment (Supplementary Table S1).

Mutant variants of DPCK were designed as follows. First, all Phe, Tyr and Trp residues were substituted by (i) Leu residues (DPCK-LH) and (ii) non-aromatic amino acids based on best preservation of thermodynamic stability (DPCK-MH) using the Hot Spot Wizard server (26). Second, all of the above amino acids plus His were substituted using the same logic, producing DPCK-L and DPCK-M variants respectively. DPCK-LH/MH and DPCK-L/M variants thus have 10% and 11% of the total protein sequence substituted, respectively. Synthetic genes of all these variants were subcloned and expressed in *E. coli* with a C-terminal histidine tag using standard protocols (see Methods for details). Upon preliminary purification and DPCK activity characterization, only DPCK-LH and DPCK-M variants were selected for detailed characterization (Fig. 1). Intriguingly, DPCK-L mutant had a very poor expression (even after optimization attempts) in *E. coli* and both DPCK-L and DPCK-MH mutants did not have any measurable DPCK-related activity (Supplementary Table S2).

**Fig. 1.**
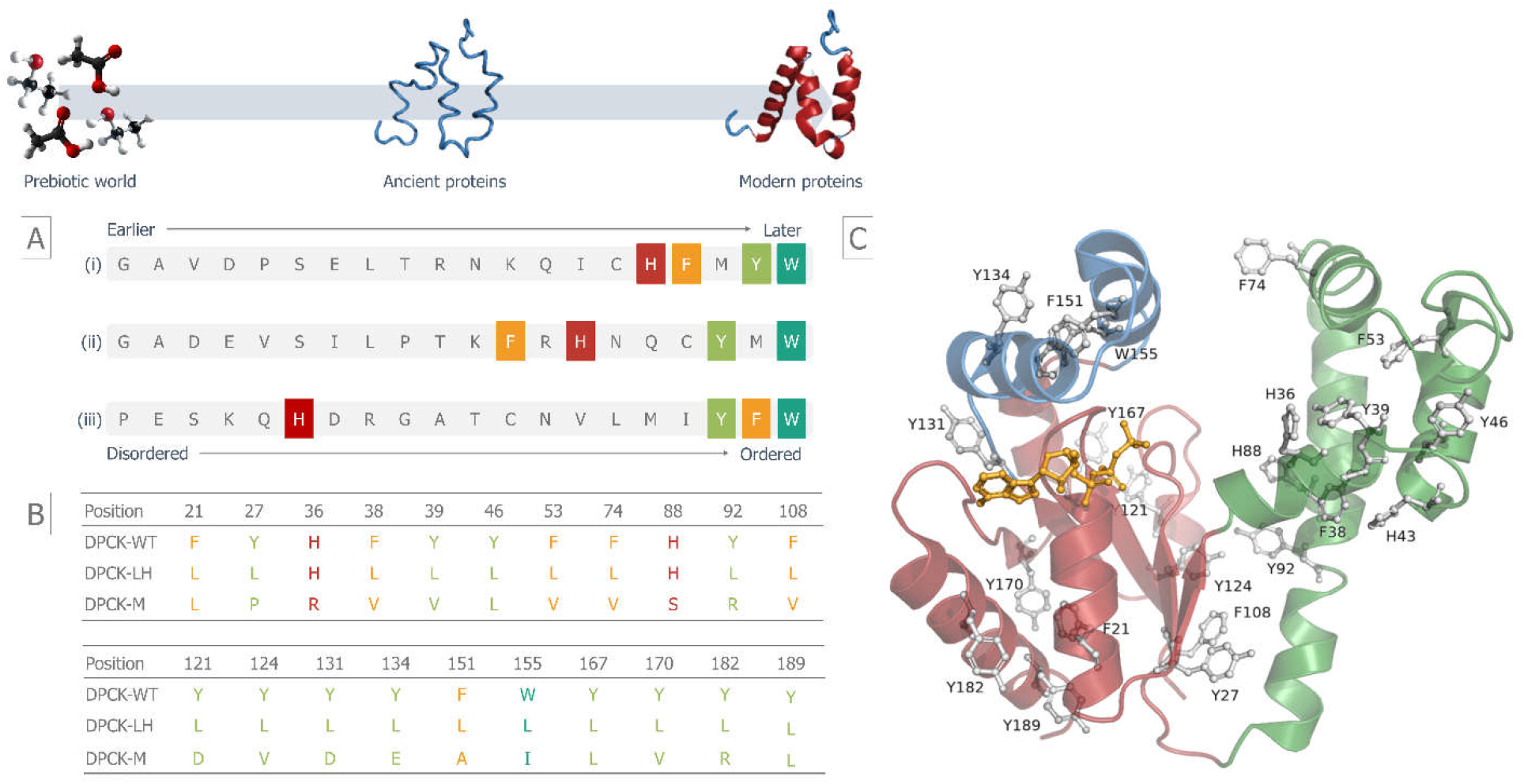
Sequence design of DPCK variants lacking aromatic amino acids. (A) The chronological order and ranking of 20 amino acids: (i) order of appearance in the genetic code derived by meta-analyses by Trifonov (9); (ii) order of appearance based on the prebiotic availability and thermodynamic stability by Higgs and Pudritz (8); (iii) ranking based on their increasing propensity to promote structure (18) (B) Aromatic amino acid content of DPCK-WT, -LH and -M variants (C) Aromatic residues highlighted in the structure of DPCK from Aquifex aeolicus (PDB ID 2IF2), with ATP molecule positioned based on structural alignment with the H. influenzae DPCK complex with ATP (PDB ID 1JJV)

DPCK-WT, -LH and -M variants were purified to homogeneity using a three-step purification protocol (Supplementary Fig. S1). Prior to further experiments, the identity, molecular weight and oligomeric status of the protein variants were tested by mass spectrometry and analytical size exclusion chromatography (Supplementary Fig. S1). All protein variants were of expected molecular weight. DPCK-WT and -LH eluted as monomers while the -M variant resembles either a dimeric or disordered monomeric form in the elution profile.

### Enzyme activity characterization

The specificity and rates of enzymatic reactions of the DPCK variants were initially characterized using a commercial kit relying on a coupling detection of ADP, one of the reaction products. In the assay, the ADP is converted to pyruvate which is then quantified by a fluorometric method. Because the assay was performed at two regimes (varying the concentration of ATP or dCoA), it was possible to observe significant differences in the reaction specificity of the variants (Fig 2).

**Fig. 2.**
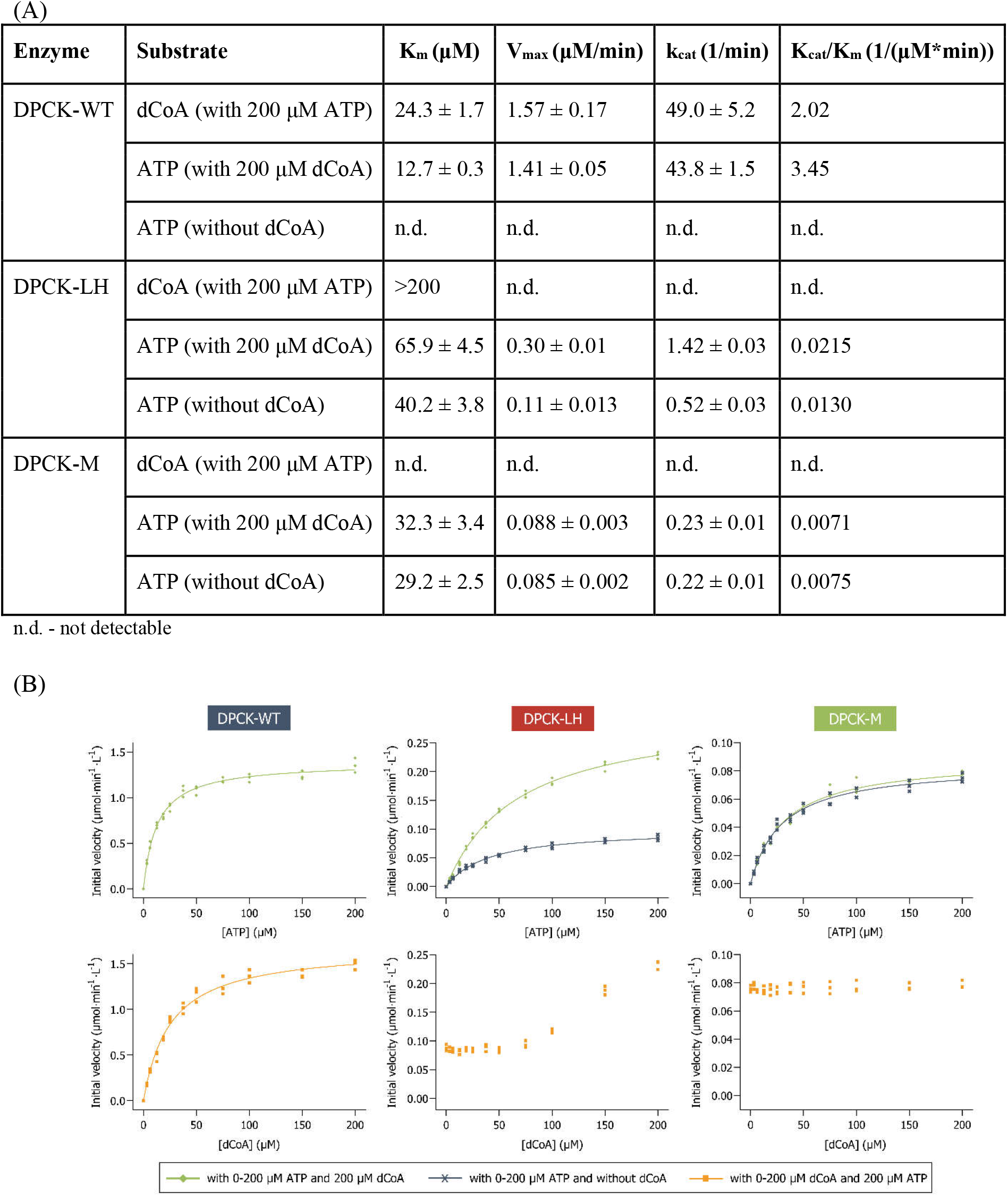
Kinetic characterization of DPCK variants. (A) Summary of catalytic efficiencies (B) Michaelis-Menten plots of DPCK proteins for initial velocity versus (TOP) ATP concentration, monitoring production of ADP; reactions were performed without and with 200 μM dCoA to estimate ATPase and phosphotransferase activities of enzymes. (BOTTOM) dCoA concentration, monitoring production of ADP. Reactions were performed in 15 mM Hepes (pH 7.4), 20 mM NaCl, 1 mM EGTA, 0.02% Tween-20, 10 mM MgCl_2_, and 0.1% bovine gamma globulin, and was initiated with ATP. The lines represent nonlinear least squares fits.

DPCK-WT has similar catalytic efficiency for both ATP and dCoA as substrates while the ATP hydrolysis activity is dependent on dCoA binding (Fig. 2). The herein measured catalytic efficiency of the reaction (2 and 3.5×10^−6^ M^−1^min^−1^ for dCoA and ATP, respectively) is similar to previously reported efficiency of DPCK from *E. histolytica* (27). In contrast, the catalytic efficiency of DPCK-LH and DPCK-M are significantly lower (2.2×10^−8^ and 0.7×10^−8^ M^−1^min^−1^for ATP, respectively), resulting in a decreased turnover number (Fig. 2 and Supplementary Fig. S2). In the case of DPCK-M, these rates are independent of varying concentrations of dCoA implying an impaired efficiency of the phosphate transfer, i.e. only phosphatase activity is observed (Fig. 2B). While both DPCK-LH and -M variants have the capacity to hydrolyze ATP in the absence of dCoA (unlike DPCK-WT), DPCK-LH has also the dCoA dependent activity (above app. 80 μM dCoA) with K_*M*_ bigger than 200 μM. This activity has been difficult to measure using the commercial kit due to the concentration range limitation. In order to confirm the identity of the reaction products and reaction specificity, the DPCK reactions were performed at a fixed substrate concentration above the DPCK-WT K_*M*_ value (where the reaction rate is less dependent or independent of substrate concentration) in order to reach sufficient substrate conversion for detection of the products using HPLC-MS analysis. This analysis detected significant CoA formation only in the reaction catalyzed by DPCK-WT and 100x lower formation was detected in the reactions catalyzed by DPCK-LH (Supplementary Fig. S2).

### Secondary and tertiary structure characterization

Using the purified proteins, their structural properties were investigated using electronic circular dichroism (ECD), NMR and limited proteolysis.

ECD spectrum of DPCK-WT (Fig. 3A) with comparable intensity of negative maxima at 209 and 225 nm and with intense positive maximum at 195 nm indicates relatively high partition of α-helical structure (∼45 %). This is confirmed by the numerical data analysis and agrees with the secondary structure assignment of the X-ray structure (PDB code 2IF2) (Supplementary Table S3). In the case of DPCK-LH, the first negative maximum is blue-shifted to 207 nm and its intensity is comparable to that of the second negative maximum at 225 nm. This together with the positive maximum at 195 nm (almost half intensity compared to CD spectrum of DPCK-WT) reveals a significant content of α-helical structure (∼40 %) together with more pronounced partition of β-sheet structure, confirmed also by the numerical data analysis (Supplementary Table S3). ECD spectrum of DPCK-M has the first negative maximum also blue-shifted up to 205 nm but this spectral band is more intense compared to the second negative maximum at 222 nm, which could imply possible formation of 3_10_-helical structure as well as enrichment of unordered structure. The overall spectral shape and mainly spectral intensity of a positive spectral band at 192 nm could be due to a relatively high portion of β-sheet structure (Supplementary Table S3).

**Fig. 3.**
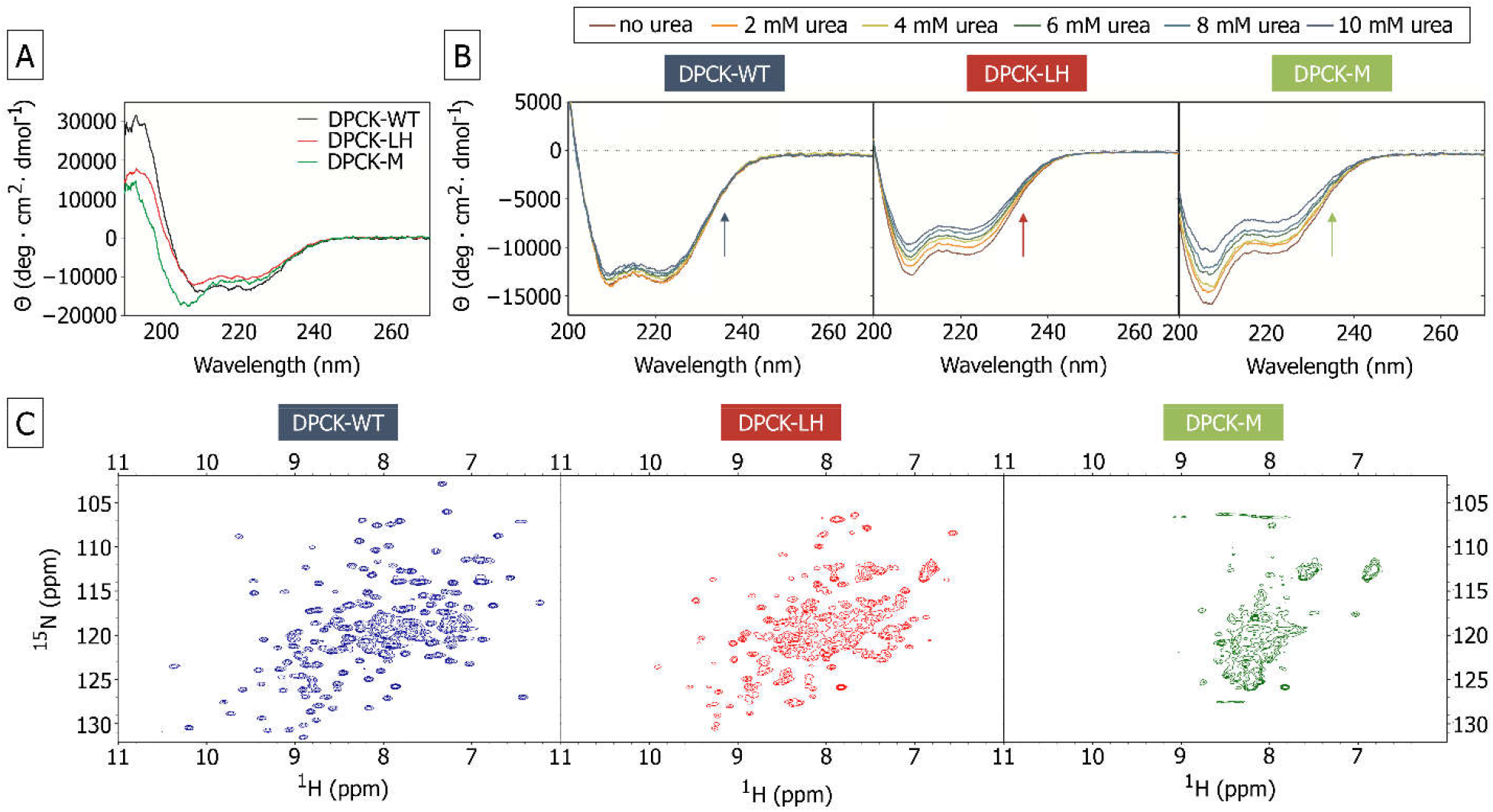
Secondary and tertiary structure characterization of DPCK variants. (A) Far-UV CD spectra of DPCK proteins. The spectra were collected in PBS buffer (11.8 mM phosphate (pH 7.6), 137 mM NaCl, 5 mM MgCl_2_, 2.7 mM KCl and 0.5 mM DTT) (B) Far-UV CD spectra of DPCK proteins measured at 0-10 mM urea. The spectra were collected in PBS buffer (11.8 mM phosphate (pH 7.6), 137 mM NaCl, 5 mM MgCl_2_, 2.7 mM KCl and 0.5 mM DTT) with addition of 0—10 mM urea (C) 2D NMR of DPCK proteins. The spectra were collected in 50 mM phosphate (pH 7.6), 280 mM NaCl, 20 mM KCl, 10 mM MgCl_2_, and 0.5 mM TCEP.

To estimate the influence of amino acid mutation on overall protein structure stability, the protein was unfolded with urea in concentration ranging from 0mM to 10mM and was further studied using CD spectroscopy. While DPCK-WT ECD spectrum remains constant upon mild urea titration (up to 10 mM), the urea titration spectra indicate a molten globular feature of in both DPCK-LH and -M variants (Fig. 3B). The DPCK-M variant, in which all aromatics were replaced, exhibited a much stronger effect than –LH variant.

Similarity of structural resemblance of DPCK-LH and DPCK-WT was further confirmed by 1D and 2D HN NMR spectra. DPCK-WT spectrum has a good signal dispersion in the -NH-region (6-9 ppm) and clear signals near 1 ppm indicative of methyl groups in the hydrophobic core, all features corresponding to a well-folded protein. While the signal of the methyl groups in the hydrophobic core is absent in the DPCK-LH variant spectrum (as expected from the removal of aromatic residues), the signal dispersion in the -NH-region implies that the -LH variant is at least partially folded, in contrast with that of the -M variant where the signal in the same region is less dispersed, implying lack of tertiary structure (Fig. 3C and Supplementary Fig. S3). Based on the analyses of N-edited 3D NOESY spectra, the following counts of α-helical peaks at 131, 57 and 14 for DPCK-WT, -LH and -M variants, respectively (Supplementary Table S3).

The tertiary structure of the proteins was additionally characterized by limited proteolysis by endoproteinase Lys-C as its cleavage site map is conserved among all studied variants (Supplementary Fig. S4). DPCK-WT shows high resistance to proteolytic digestion during the whole-time scale of the limited proteolysis experiment, reflecting its globular structure. In contrast, both mutant variants are gradually digested by Lys-C over time and the amounts of the intact DPCK-LH and DPCK-M decrease exponentially over time. While relatively large cleaved fragments with the approximate size of 15 kDa can be observed during proteolysis of DPCK-LH, no large cleavage fragments are detected during proteolysis of DPCK-M, an indication of its loose or absent tertiary structure (Fig. 4).

**Fig. 4.**
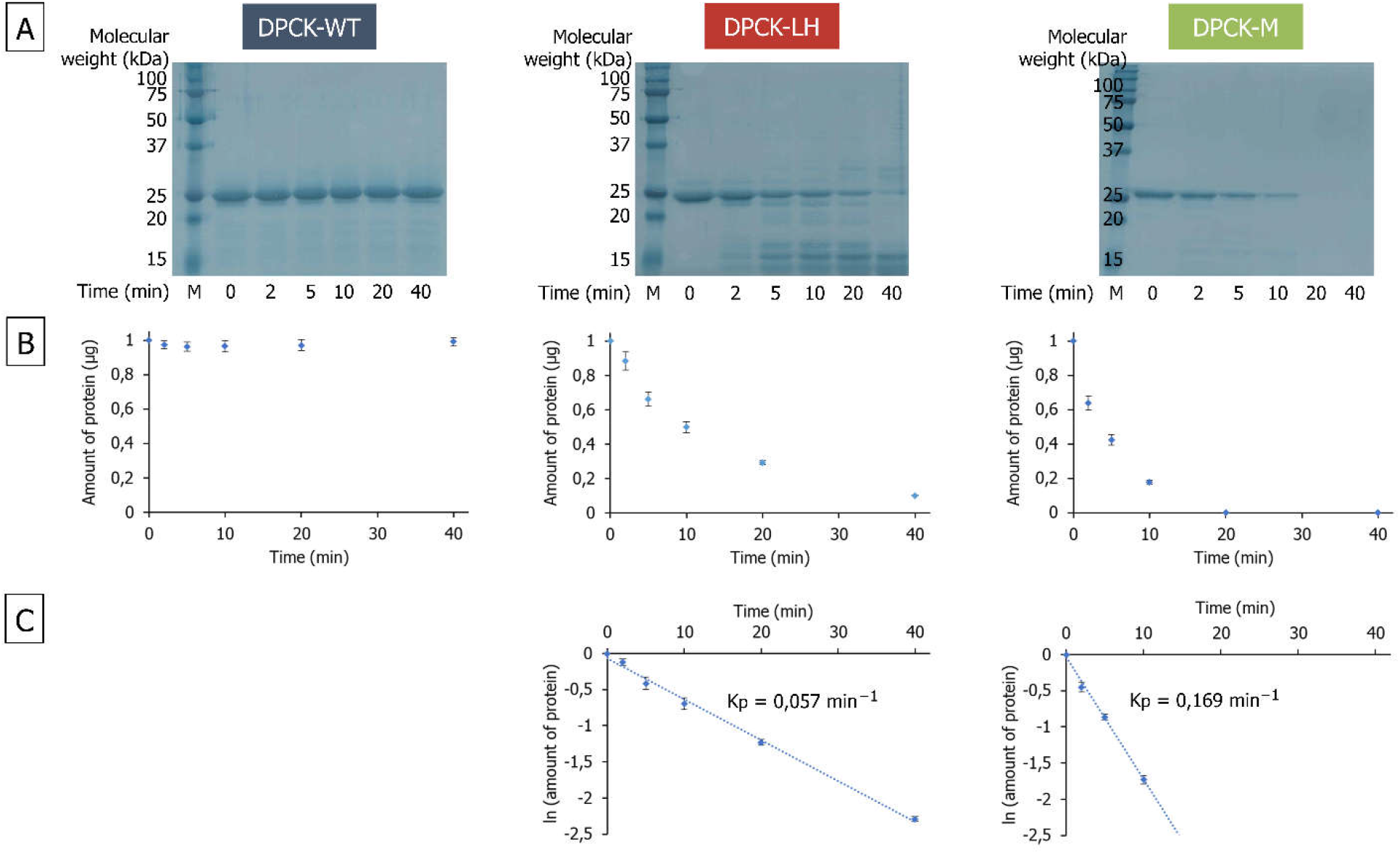
Limited proteolysis of DPCK proteins. (A) 14% SDS-polyacrylamide gels visualized by imidazole-zinc staining after SDS-PAGE with the protein samples exposed to Lys-C endoproteinase for different times. (B) Graphs representing the amount of the proteins remaining at each time point. (C) Determination of proteolysis rate constants (K_p_) assuming pseudo-first order of proteolytic reactions.

In summary, DPCK-LH variant (which has all the aromatic amino acids substituted by leucine) shows relatively high conservation of secondary structure but a loose tertiary structure (probably of molten globular nature) when compared with the -WT protein. On the other hand, both secondary and tertiary structures of DPCK-M variant are severely impaired, in which all histidines were substituted in addition to aromatics.

### Structural characterization of ATP binding

For an efficient phosphorylation reaction, the ATP gamma-phosphate has to be protected from a nucleophilic attack by water molecules. DPCK active site must therefore be shielded from water once ATP molecule is bound. Similar to other kinases, this conformational change of DPCK, induced by ATP binding, has been described previously (28).

To study the structural changes of DPCK variants upon ATP binding, 2D HN NMR spectra were collected in response to ATP titration (see Supplementary Fig S5). While the NMR spectra of the DPCK-M variant are of generally low quality, which is probably caused by complex dynamics on the millisecond time-scale making the protein signals invisible for NMR spectroscopy, the spectra of both DPCK-WT and -LH variants show expected perturbations upon ATP titration. For DPCK-WT we observe typical examples of slow exchange behavior, where only free and bound forms are observed with peak intensity proportional to the population. Interestingly, for the DPCK-LH variant, we typically observed examples of fast exchange with only a single peak visible at a given protein:ATP ratio, although examples of slow exchange are observed as well (Fig. 5A). This suggests that compared to the WT:ATP interaction, an additional process occurs during DPCK-LH titration with ATP.

**Fig. 5.**
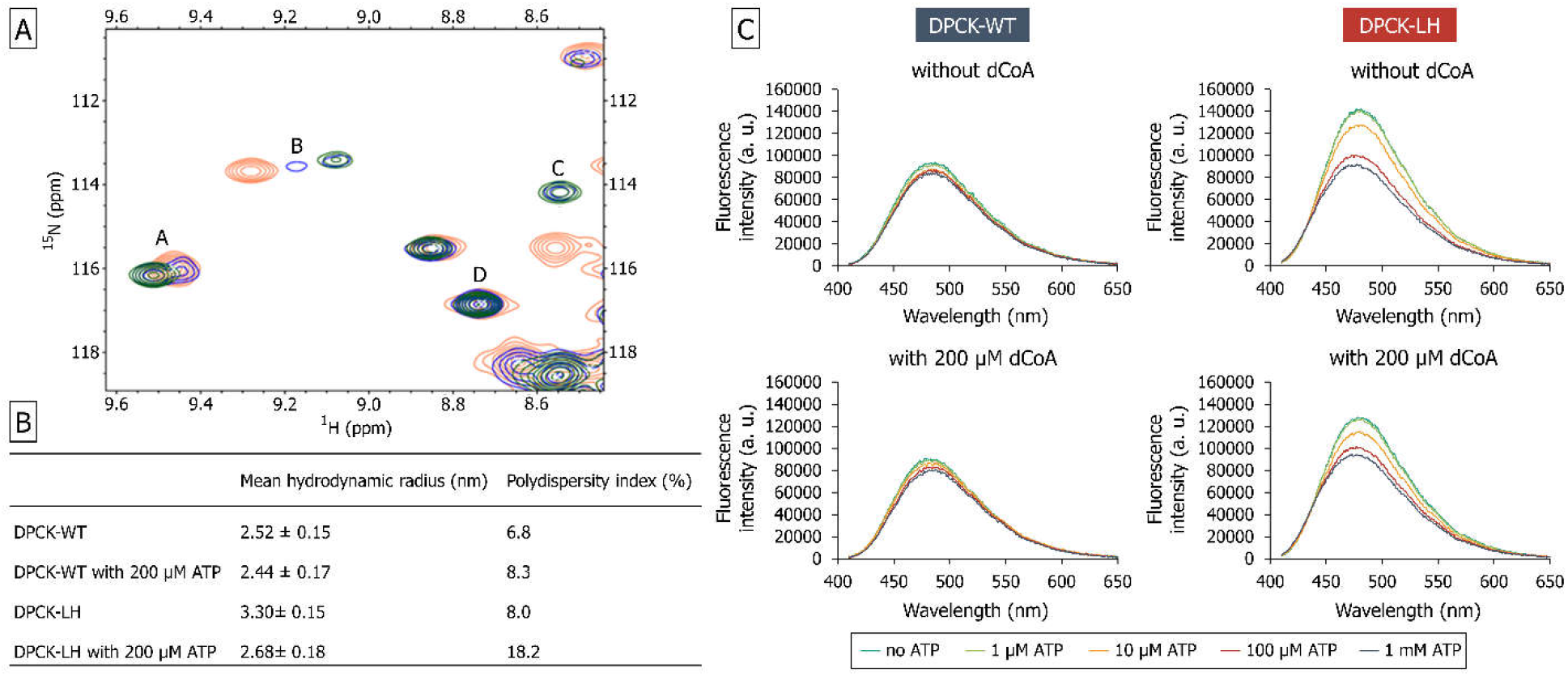
Structural Characterization of DPCK-WT and -LH upon substrate binding. (A) An exemplary close-up of DPCK-LH 2D NH NMR spectra induced by ATP binding (red – free protein (100 μM), blue – 300 μM ATP, green – 1000 μM ATP). (B) Mean hydrodynamic radius of DPCK-WT and -LH variants with and without 200 μM ATP measured by dynamic light scattering. (C) The steady-state fluorescence spectra of ANS binding at excitation wavelength 380 nm. The spectra were measured at different concentrations of ATP (with and without 200 μM dCoA), and each spectrum is the average of three individual scans. The fluorescence was recorded between 410 – 650 nm after exciting the protein solution at 380 nm.

To further investigate this intriguing observation, DPCK-WT and -LH (i.e. those variants that are capable of phosphotransferase activity that requires a hydrophobic core) structural response to substrate binding was tested using dynamic light scattering and ANS titration. The steady-state fluorescence measurements support the molten globule nature of DPCK-LH variant since it shows higher fluorescence intensity values in comparison with DPCK-WT, resulting from the high affinity of ANS to the exposed hydrophobic core of molten globular intermediates (29). While the fluorescence intensity decreases for both variants upon substrate binding, this change is significantly more dramatic for DPCK-LH (Fig. 5C). ATP (out of the two substrates) has a remarkable effect on additional folding of DPCK-LH protein, explaining its ability to perform the phosphotransferase activity despite its molten globular nature in the free state. Both 2D HN NMR and ANS titration observations were further supported by DLS measurements where the mean hydrodynamic radius of DPCK-LH was recorded to be reduced by ∼20% and reached that of DPCK-WT value upon ATP addition (Fig. 5B).

## Discussion

Aromatic residues are essential for formation of a stable hydrophobic core of extant proteins (20). At the same time, tight protein folding is frequently required for enzyme catalysis even though most enzymes undergo dynamic structural changes during the process. With aromatics being apparently the latest addition to the amino acid alphabet, how specific protein catalysis could be achieved in their absence remains unclear. The work reported here sheds some light on this problem.

To examine the contribution of the aromatic amino acids to enzyme catalysis, we performed a detailed analysis of two aromatics-less mutants of the *Aquifex aeolicus* DPCK where (i) all Phe, Tyr and Trp residues were substituted by Leu residues (DPCK-LH), and (ii) all Phe, Tyr, Trp and His were substituted by non-aromatic amino acids based on predicted preservation of thermodynamic stability (DPCK-M). According to the Trifonov’s chronology of amino acid incorporation into genetic coding, His was among the last amino acids added (9). On the other hand, according to the order-disorder propensity scale, His is among the most disorder-promoting amino acids, likely due to its significant positive charge and the two hydrogen-bonding nitrogen groups that could promote structural instability by bond switching (19). In contrast to the tendency of His to induce disorder, in the presence of divalent cations, especial zinc, His often plays in role in inducing structure due to its metal ion coordination.

The wild type variant of DPCK performs phosphotransferase activity from ATP to dCoA where dCoA acts as the leading substrate and ATP does not get hydrolysed in its absence (30). The unliganded DPCK is a well folded protein with high α-helical content, its domain movements upon ATP binding play a crucial role during catalysis (28). None of the aromatic amino acid residues has been reported essential for the ligand binding and catalysis (30).

Interestingly, the DPCK-M variant (selected for best predicted preservation of thermodynamic stability) had a more impaired structural integrity than DPCK-LH. This may be either the consequence of the specific substitutions or the indispensability of the DPCK’s His residues. Given the speculative role of His for hydrogen bond switching, it would be interesting to determine whether His plays a role in facilitating the domain movements needed for catalysis. In contrast, CD and NMR measurements of the DPCK-LH variant showed a similar content of secondary structure to the wild type protein but limited proteolysis, CD urea titration and 2D NMR all imply its molten globule tertiary conformation. DPCK-M variant has no measurable phosphotransferase activity while both of the mutant variants are able to hydrolyse ATP even in the absence of dCoA. This is likely due to the loss of structural orchestration of the catalytic events and demonstrates that some activities can be performed even in the absence of a firm hydrophobic core. However, this is probably untrue for the phosphotransferase activity where the gamma-phosphate has to be protected from a nucleophilic attack by water molecules in order to be efficiently transferred to the desired substrate. DPCK-LH variant is still able to perform this activity although with significantly lower efficiency (∼100×) compared with the wild type protein. Similar to the wild type protein, slow-exchange behavior is observed in the NMR spectra upon ATP titration, representing the ATP-induced change in structural conformation. However, the DPCK-LH variant undergoes significant additional folding upon ATP binding, explaining its ability to perform the phosphotransferase reaction. This ligand-induced folding scenario is in agreement with previously reported engineered molten globular enzymes (31-33). In a well-structured, lock and key type enzyme, most of the binding energy translates into substrate distortion so large rate accelerations can be achieved. However, for molten globular enzymes such as the one described here, some of the binding energy is needed to induce protein folding so one expects a reduced rate acceleration as we have found here.

The study of the engineered molten globular enzyme (32) includes the hypothesis that modern enzymes evolved from molten globular precursors. If the earliest cells indeed existed without aromatics, then the data in this paper adds weight to this evolutionary scenario of molten globular polypeptide → molten globular enzyme → modern enzyme, where the last step is enabled by the expansion of the genetic code to include the aromatic residues. In this view, the evolutionary advantage of adding the aromatic amino acids was to enable the formation of tightly structured proteins with an accompanying high performance catalysis.

To further test this idea, work in progress is to use bioinformatics tools to carry out disorder prediction with VSL2B on DPCK and the other identified ancient enzymes with their modern sequences and with their aromatics replaced by Leu. All of the ancient enzymes so far tested are predicted to be structured with the aromatics and disordered without these residues. The next step will be to apply additional bioinformatic tools that distinguish molten globules from other types of disorder (34)

In summary, we report an enzyme without aromatic amino acids that is still capable of a specific, hydrophobic pocket dependent catalysis. This enzyme is rich in secondary structure but exhibits a molten globule conformation in an unliganded form. Our study provides evidence that a tightly packed protein environment can form upon its ligand binding. This phenomenon could be relevant in the early stages of enzyme catalysis before the fixation of the contemporary amino acid alphabet.

## Methods

### Plasmid preparation

DPCK genes for DPCK-WT, DPCK-LH, -L, -MH and -M were amplified by PCR using Pfu-X DNA polymerase (Jena Bioscience, Germany) according to the following program: an initial denaturation at 95 °C for 2 min; followed by the 32 cycles of denaturation at 95 °C for 30 sec; annealing at 56 °C for 30 sec; and elongation at 68 °C for 30 sec; and a final extension at 68 °C for 2 min. The PCR amplification for all three genes was performed with the same set of primers: forward, 5’-AAAAA<u>CATATG</u>AAACGTATCGGTCTGACC-3’, and reverse, 5’-AAAAA<u>CTCGAG</u>TTCCAGCGGGTCACGG-3’. The PCR fragments were digested with *XhoI* (New England BioLabs, USA) and *NdeI* (New England BioLabs, USA), purified with Monarch PCR & DNA Cleanup Kit (New England BioLabs, USA) and cloned into PET-24a (+) C-terminal-histidine-tag vector (Novagen, Germany), which was digested by *XhoI* and *NdeI* and dephosphorylated by Antarctic Phosphatase (New England BioLabs, USA) prior to ligation.

The plasmids were introduced into One Shot TOP10 Chemically Competent *E. coli* cells (Thermo Fisher Scientific, USA) by heat shock protocol at 42 °C for 60 sec, and the cells were grown overnight at 37 °C on LB agar plates containing 50 μg/ml of kanamycin (Sigma Aldrich, USA). A single colony was selected, cells were grown overnight at 37 °C in 5 ml of LB Broth (Sigma Aldrich, USA) supplemented with 50 μg/ml of kanamycin (Sigma Aldrich, USA) and plasmid DNA was isolated and analyzed by Sanger sequencing.

### Protein expression and purification

Isolated plasmids were introduced into *E. Coli* BL21 (DE3) cells (Thermo Fisher Scientific, USA), and the cells were grown overnight at 37 °C in 5 ml of LB Broth (Sigma Aldrich, USA) in the presence of 50 μg/ml of kanamycin. The overnight cultures were used to inoculate 500 ml of fresh medium, and the culture was propagated at 37 °C at 220 rpm shaking. When OD_600_ reached 0.7-0.8, isopropyl β-D-thiogalactopyranoside (IPTG, Sigma Aldrich, USA) was added to final concentration of 0.5 mM and the cultivation was continued for 4 h at 37 °C. The cells were harvested by centrifugation at 3000× g for 20 min at 4 °C. The cell pellets were resuspended in 15 ml of lysis buffer (20 mM Tris (pH 8.0), 20 mM NaCl, and 1 mM of β-mercaptoethanol) with one tablet of EASYpack protease inhibitor cocktail (Sigma Aldrich, USA), incubated with 50 μg/ml of lysozyme and 6 U of RNase-free DNase I (Jena Bioscience, Germany) at room temperature for 30 min, sonicated on ice at 1.5 W (18 cycles, 10 sec on, 20 sec off) and centrifuged at 35000× g for 30 min at 4 °C. After, Tween-20 (Sigma Aldrich, USA) was added to supernatants to the final concentration of 0.1 % (v/v), and the crude lysates were applied to 5 ml HiTrap Capto Q column (GE Healthcare Life Sciences, USA) equilibrated with 5 volumes of buffer A (20 mM Tris (pH 8.0), 20 mM NaCl, 1 mM beta-mercaptoethanol and 0.1 % (v/v) Tween-20). Then, the DPCK proteins were eluted with 0-50% gradient of buffer B (20 mM Tris (pH 8.0), 1 M NaCl, 1 mM beta-mercaptoethanol and 0.1 % (v/v) Tween-20), and fractions from 15 % to 35 % of buffer B were collected and applied to 5ml HisTrap HP column (GE Healthcare Life Sciences, USA) equilibrated with 5 volumes of buffer C (20 mM Tris (pH 7.6), 500 mM NaCl, 10 mM imidazole, 1 mM beta-mercaptoethanol and 0.1 % (v/v) Tween-20). The column was washed with 3 % of buffer D (20 mM Tris (pH 7.6), 500 mM NaCl, 500 mM imidazole, 1 mM beta-mercaptoethanol and 0.1 % (v/v) Tween-20) to remove unbound proteins, and the DPCK proteins were eluted with 0 — 50 % gradient of buffer D. Fractions from 20 % to 30 % of buffer D were collected, concentrated up to 0.5 ml by centrifugation using 4 ml Amicon Ultra centrifugal units (MWCO 10 000, Millipore, USA) and applied to Superdex 75 10/300 GL column (GE Healthcare Life Sciences, USA) equilibrated with 2 column volumes of buffer E (50 mM Tris (pH 7.6), 500 mM NaCl, 20 mM KCl, 10 mM MgCl_2_ and 0.5 mM DTT). The DPCK variants were eluted as single peaks with approximate sizes of 29 kDa (DPCK-WT), 33 kDa (DPCK-LH) and 55 kDa (DPCK-M). Molecular weights were estimated using Gel filtration low molecular weight calibration kit (GE Healthcare Life Sciences, USA). After the confirmation of proteins integrity and purity by SDS-PAGE analysis on 14% acrylamide gel, the purified proteins were concentrated up to 10 mg/ml concentration and aliquoted. The aliquots were flash frozen in liquid nitrogen and stored at –80 °C.

### Basic biophysical characterization

The identities and molecular weights of purified proteins were confirmed by mass spectrometry using UltrafleXtreme MALDI-TOF/TOF mass spectrometer (Bruker, Germany) according to the standard procedure. Protein concentrations were determined by amino acid analysis using a Biochrom 30+ Series Amino Acid Analyser (Biochrom, UK).

The size distribution of protein samples was characterized using dynamic light scattering (DLS) technique. Protein samples were diluted in PBS buffer (11.8 mM phosphate buffer (pH 7.6), 137 mM NaCl, 5 mM MgCl_2_, 2.7 mM KCl and 0.5 mM DTT) to the final concentration of 0.5 mg/ml and centrifuged at 25000× g for 30 min at 4 °C. In order toremove dust particles, samples were filtered using 0.22 μm Ultrafree-MC centrifugation filters (Millipore, USA). The DLS measurements were performed in a quartz glass cuvette (light path 10 mm) at 18 °C using a laser spectroscatter-201 system (RiNA GmbH Berlin, Germany). A series of 35 measurements with a sampling time of 30 sec and a wait time of 1 sec was conducted for each sample. A diode laser of wavelength 685 nm and an optical power of 30 mW was used as the source. The scattered light was collected at a fixed scattering angle of 90º, and the autocorrelation functions were analyzed with the program CONTIN to obtain hydrodynamic radius distributions. DLS measurements were performed for protein samples in the presence of 200 μM ATP to estimate the effect of ATP binding on the hydrodynamic radius of proteins.

### Enzyme assays

DPCK activities of recombinant proteins were measured by a coupling assay using ADP Quest Assay kit (Eurofins DiscoverX, USA) according to the manufacturer’s instructions. Enzymatic assays were carried out using 80 ng (32 nM) of DPCK-WT, 500 ng (214 nM) of DPCK-LH and 900 ng (386 nM) of DPCK-M and two kind of substrates, 0–200 μM for dephospho-CoA (dCoA) at 200 μM ATP and 0–200 μM for ATP without and with 200 μM dCoA to estimate ATPase and phosphotransferase activities of enzymes. All reactions were performed in assay buffer containing 15 mM Hepes (pH 7.4), 20 mM NaCl, 1 mM EGTA, 0.02 % Tween-20, 10 mM MgCl_2_, and 0.1 % bovine gamma globulin in 96-well black microplate with 40 μl total volume. After 20 μl of reagent A and 40 μl of reagent B were added, the plates were heated at 37 °C for 10 min, and the reactions were started by adding ATP. The fluorescent intensity signal was measured at 37 °C in kinetic mode with 2 min intervals using CLARIO star microplate reader (BMG LABTECH, Germany) at excitation/emission wavelengths of 530/590 nm. The kinetic parameters were calculated using the non-linear regression function using the single saturating concentrations of substrates. Substrate conversion did not exceed 10 %. The experiments were repeated three times, and kinetic values are presented as the means ± SE.

HPLC-MS analysis was used for comparative detection of the reaction analytes. For this purpose, 100 μl of reaction mixtures were prepared by mixing 1 μg (0,42 μM) of protein, 100 μM dCoA and 100 μM ATP in 25 mM NH_4_HCO_3_ (pH 7.6), 300 mM NaCl, 20 mM KCl and 10 mM MgCl_2_. The reaction mixture was incubated at 37 °C for 1 hour, then, reaction was stopped by adding 100 μl of acetonitrile (Sigma Aldrich, USA). Precipitated recombinant protein was separated by centrifugation at 20000x g at 4 °C for 20 min.

The reaction samples were analyzed using the Dionex Ultimate 3000RS HPLC equipped with TSQ Quantiva MS detector (Thermo Fisher Scientific, USA). The ESI source was used for ionization in a positive mode. The HPLC solvent system consisted of 10 mM (NH_4_)_2_CO_3_ (pH 9.3) (A) and 97% acetonitrile (B). 1 μl of sample was injected in 50 % B and the analysis was performed using the gradient of 15 % A and 85 % B for 3.5 min followed by an increase to 75 % A and 25 % B over 11.5 min and its continuation for 10 min with the SeQuant® ZIC®-*p*HILIC column (5 µm, 150 mm x 2.1 mm, Merck, USA), at a flow rate of 0.13 ml/min.

### Circular dichroism spectroscopy

ECD spectra were collected using a Jasco 1500 spectrometer (JASCO, Japan) in the 195 —280 nm spectral range using a 0.01 cm cylindrical quartz cell. The experimental setup was as follows: 0.05 nm step resolution, 5 nm/min scanning speed, 16 s response time, 1 nm spectral band width and 2 accumulations. After baseline correction, the spectra were expressed as molar ellipticity per residue θ (deg·cm^2^·dmol^−1^). The protein samples were diluted in PBS buffer (11.8 mM phosphate (pH 7.6), 137 mM NaCl, 5 mM MgCl_2_, 2.7 mM KCl and 0.5 mM DTT) with addition of 0 — 10 mM urea. The blank spectrum of an aqueous buffer (with or without urea in a corresponding concentration) was used to correct the observed spectrum of the sample.

### Limited proteolysis

Kinetic studies on specific proteolytic cleavage by Lys-C endoproteinase were performed as follows. First, recombinant proteins were diluted in Lys-C cleavage buffer (25 mM Tris (pH 8.0), 300 mM NaCl, 1 mM EDTA, and 0.5 mM TCEP) to the final concentration of 1 mg/ml, and then reaction mixtures for proteolytic digestion were prepared by mixing 7 μl of 1 mg/ml recombinant protein and 56 μl of Lys-C cleavage buffer. After incubation at 37 °C for 10 min proteolytic cleavage was initiated by adding 7 μl of 5 ng/μl Lys-C endoproteinase. After 0, 2, 5, 10, 20 and 40 min of incubation at 37 °C 10 μl of the reaction mixture was taken out, and Lys-C was inactivated by adding 2 μl of 6x SDS-PAGE sample buffer (375 mM Tris-HCl (pH 6.8), 9 % SDS, 50 % glycerol, 9 % beta-mercaptoethanol and 0.03% bromophenol blue) followed by heating at 95 °C for 10 min. All samples then were subjected to SDS-PAGE.

For quantitative evaluation of limited proteolysis the rate constants of proteolysis were determined by monitoring the disappearance of an intact protein in a proteolysis reaction by SDS-PAGE. The areas of the bands corresponding to the intact proteins were estimated from the gels using the ImageJ program and then expressed as the amount of protein remaining after each time point. Assuming the pseudo-first order kinetics, the natural logarithms of the intact protein amounts were plotted against the time, and the plots were fitted with a first-order rate equation.

### Steady-state ANS fluorescence

Steady-state fluorescence measurements were performed using CLARIO star microplate reader (BMG LABTECH, Germany). Protein samples were diluted to 2 μM with ANS buffer (100 mM Tris, 300 mM NaCl, 20 mM KCl, 10 mM MgCl_2_ and 0.5 mM DTT) and incubated with 0; 1; 10; 100 and 1000 μM of adenosine 5′-[γ-thio]triphosphate tetralithium salt (ATP-γ-S, Sigma Aldrich, USA) at 20 °C for 30 min. After incubation, 8-anilino-1-naphthalenesulfonic acid ammonium salt (ANS, Sigma Aldrich, USA) was added to the reaction mixtures to the final concentration of 400 μM, and the reaction mixtures were incubated for additional 5 min. The final volume of each reaction mixture was 50 μl. The ANS fluorescence was excited at 380 nm, and emission spectra were recorded between 410 and 650 nm. To estimate the conformational changes induced upon dCoA binding the fluorescence intensity measurements were performed for protein samples in the presence of 200 μM dCoA. All measurements were performed in triplicates and then averaged to yield steady-state fluorescence spectra of ANS binding.

### NMR spectroscopy

NMR spectra were obtained using the Bruker© Avance HD III 850 MHz instrument, equipped with triple-resonance cryo-probe. Sample volume was 0.16 ml in 3 mm NMR tubes, in a 50 mM phosphate (pH 7.6) buffer, 280 mM NaCl, 10 mM MgCl_2_, 20 mM KCl and 0.5 mM TCEP. Protein concentration was 150 μM for 3D ^15^N/^1^H NOESY-HSQC spectra, 30 μM for DPCK-WT ATP titration, and 100 μM for both aromatic amino acid-lacking mutants ATP titration. All proteins used in the study were ^15^N labeled. ATP titrations were followed using a series of standard 1D and 2D HN correlation spectra.

## Author Contributions

MM and JM planned and performed the experiment and took part in writing the paper; VGG, VT and JV contributed significantly to the conception of the project, planned the experiments and helped with data analyses. AB, PS and LB performed the HPLC-MS, NMR and CD experiments respectively and helped with their analyses. AKD and KH planned the experiments, analyzed the data and wrote the paper.

## Acknowledgement

We would like to thank Dr. Radko Souček for help with the amino acid analysis and Dr. Tereza Ormsby, Dr. Rozálie Hexnerová and Dr. Václav Veverka for helpful suggestions. This work was supported by the Czech Science Foundation (GAČR) grant number 17-10438Y, project SVV260572/2020 (VT and MM) and by the project BIOCEV (CZ.1.05/1.1.00/02.0109), from the European Regional Development Fund. We would also like to acknowledge the facility and support of the CMS-Biocev (“Biophysical techniques”) supported by MEYS ČR (LM2018127).

