## Supplementary data for "Enzyme catalysis prior to aromatic residues: reverse engineering of a dephosphoCoA kinase"

* shared authorship

**Supplementary information**

***Supplementary Figures***

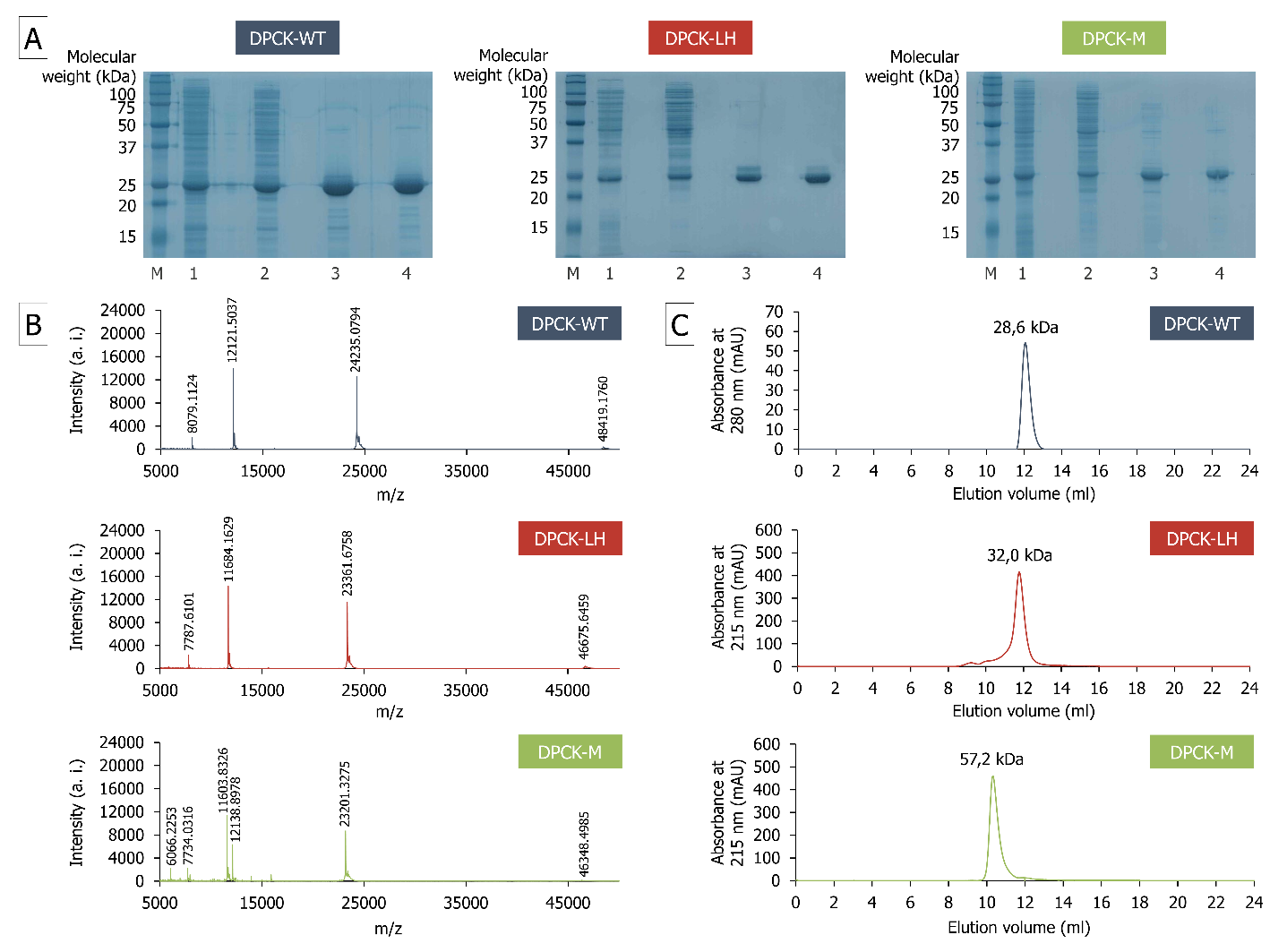

**Supplementary Fig. S1**: Purification and quality control of the studied protein variants

*(A) 14% SDS-polyacrylamide gels summarizing DPCK proteins purification course: lane M - Precision Plus Protein Dual Xtra Prestained Protein Standards, lane 1 - cell lysate, lane 2 - protein sample after anion exchange chromatography, lane 3 - protein sample after immobilized metal affinity chromatography, lane 4 - protein sample after size exclusion chromatography.*

*(B) Confirmation of the molecular weight by MALDI-TOF. The theoretically estimated molecular weights are 24,24 (DPCK-WT), 23,36 (DPCK-LH) and 23,20 (DPCK-M) kDa. The lower m/z peaks represent multiply charged species of the same proteins.*

*(C) Characterization by analytical size exclusion chromatography. The chromatograms were obtained for 50 μg of proteins in PBS buffer (11.8 mM phosphate (pH 7.6), 137 mM NaCl, 5 mM MgCl_2_, 2.7 mM KCl and 0.5 mM DTT).*

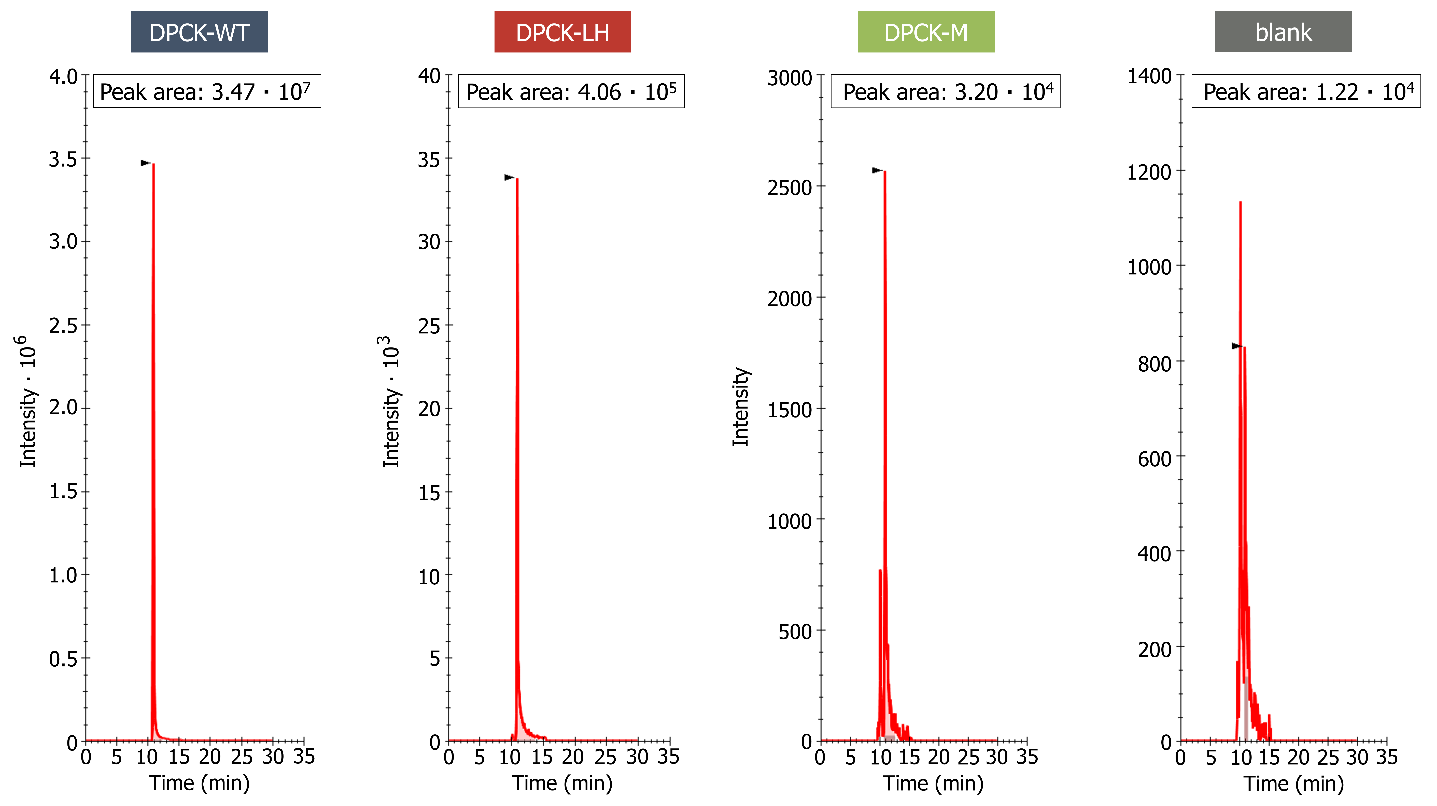

**Supplementary Fig. S2:** Detection of CoA formation by HPLC-MS at zero-order enzyme kinetics (for DPCK-WT). The reaction with both substrates at 100 μM was incubated for 1h and the CoA formation was analyzed by HPLC on the ZIC-HILIC column in acetonitrile gradient.

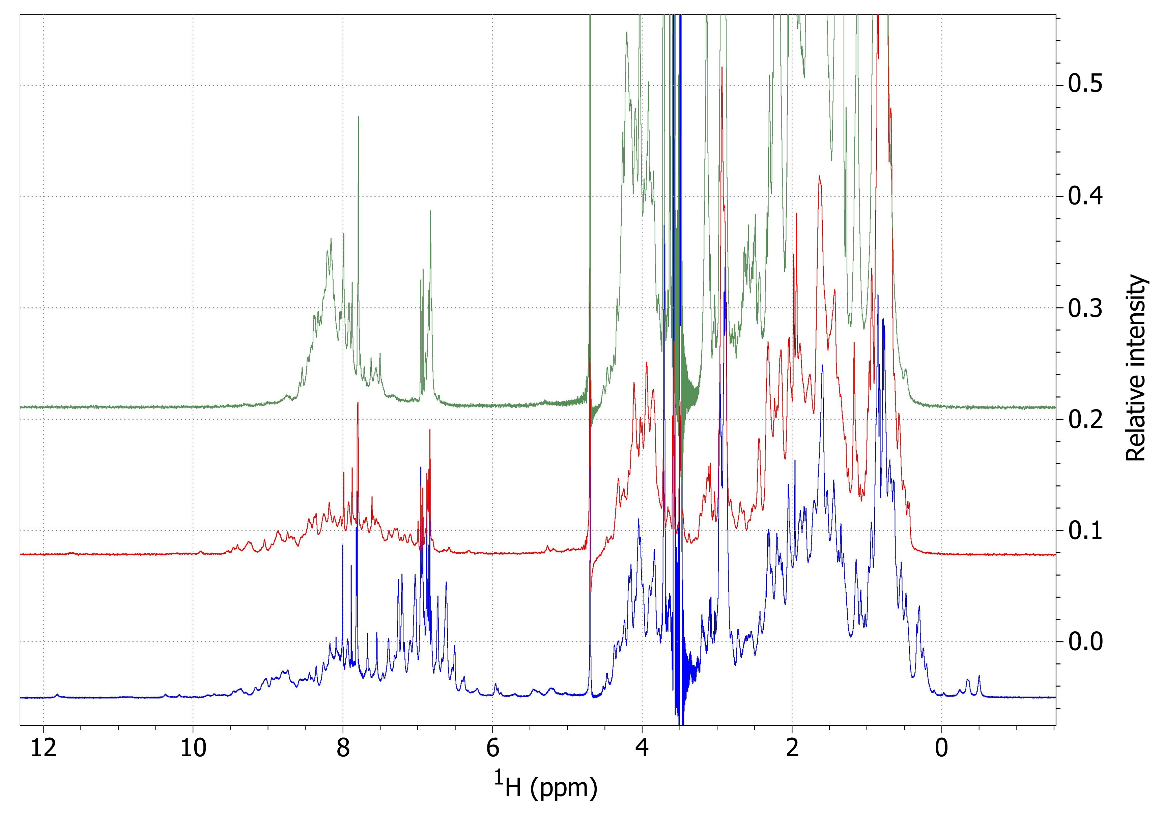

**Supplementary Fig. S3:** 1D NMR spectra: DPCK-WT - blue, DPCK-LH - red, DPCK-M - green of DPCK variants. 1D NMR spectra were collected in 50 mM phosphate (pH 7.6), 280 mM NaCl, 20 mM KCl, 10 mM MgCl_2_, and 0.5 mM TCEP.

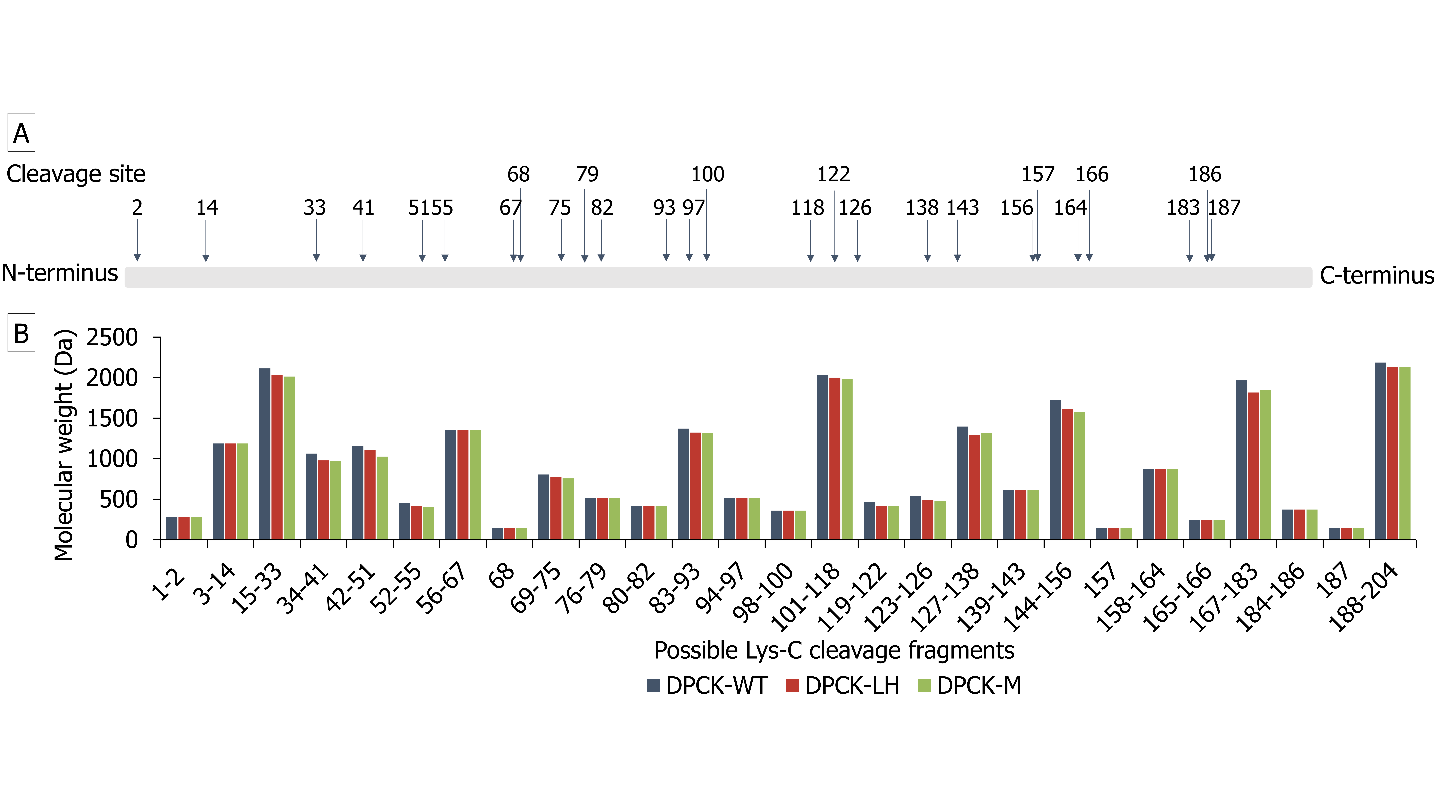

**Supplementary Fig. S4**: Lys-C limited proteolysis characterization (A) Map of possible Lys-C cleavage sites and (B) Molecular weights of possible Lys-C cleavage fragments calculated using PeptideCutter ExPASy tool

(A)

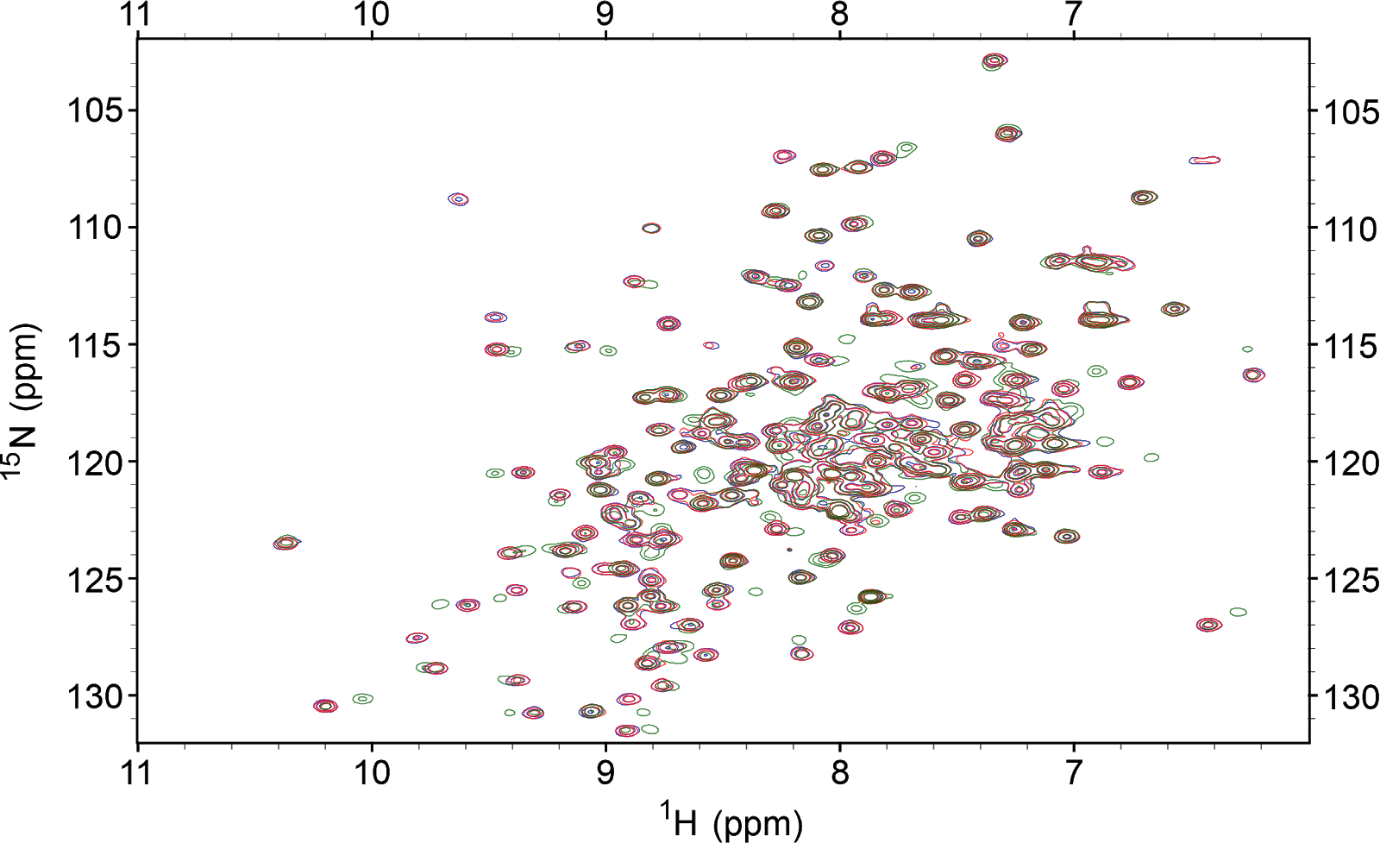

(B)

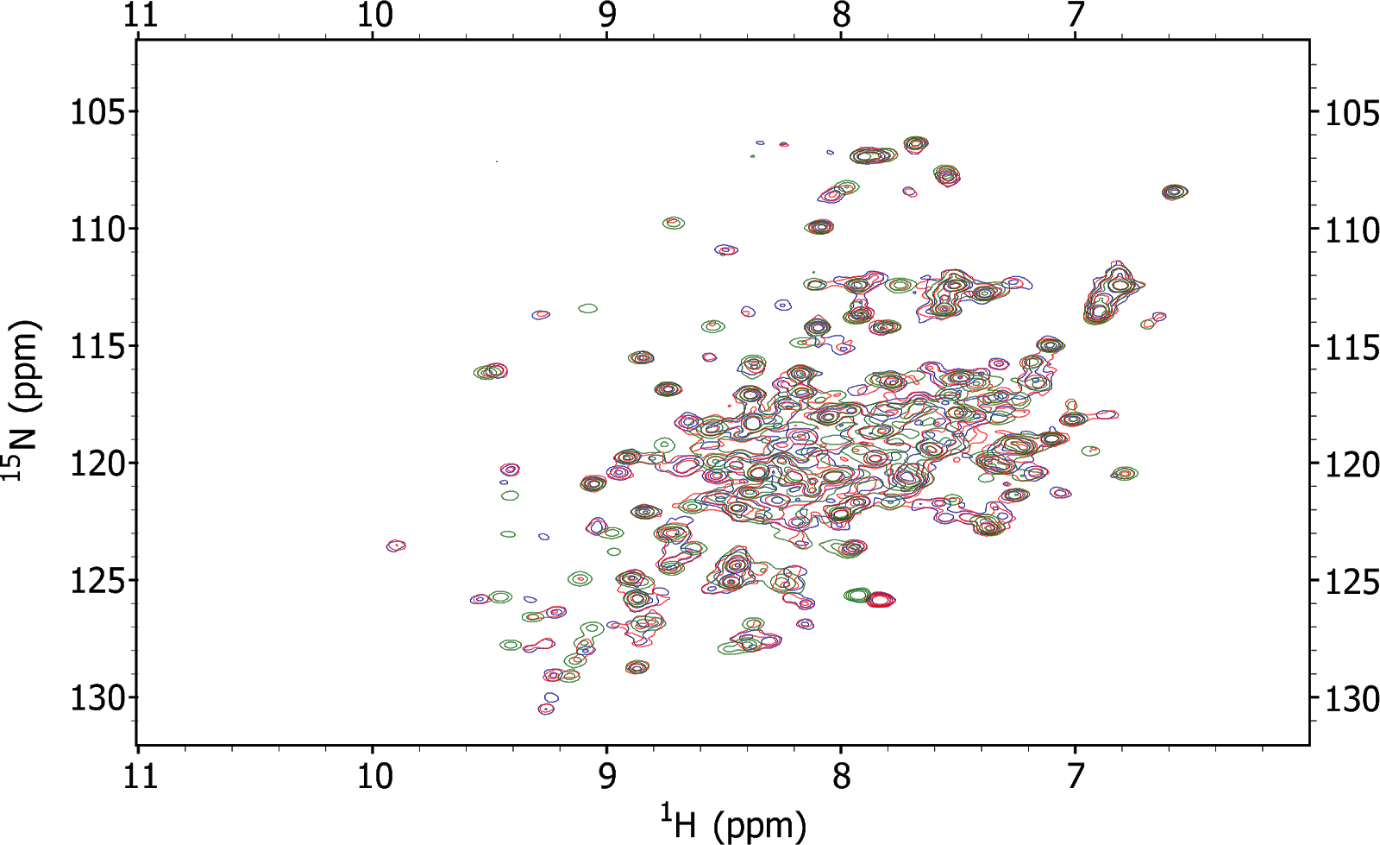

(C)

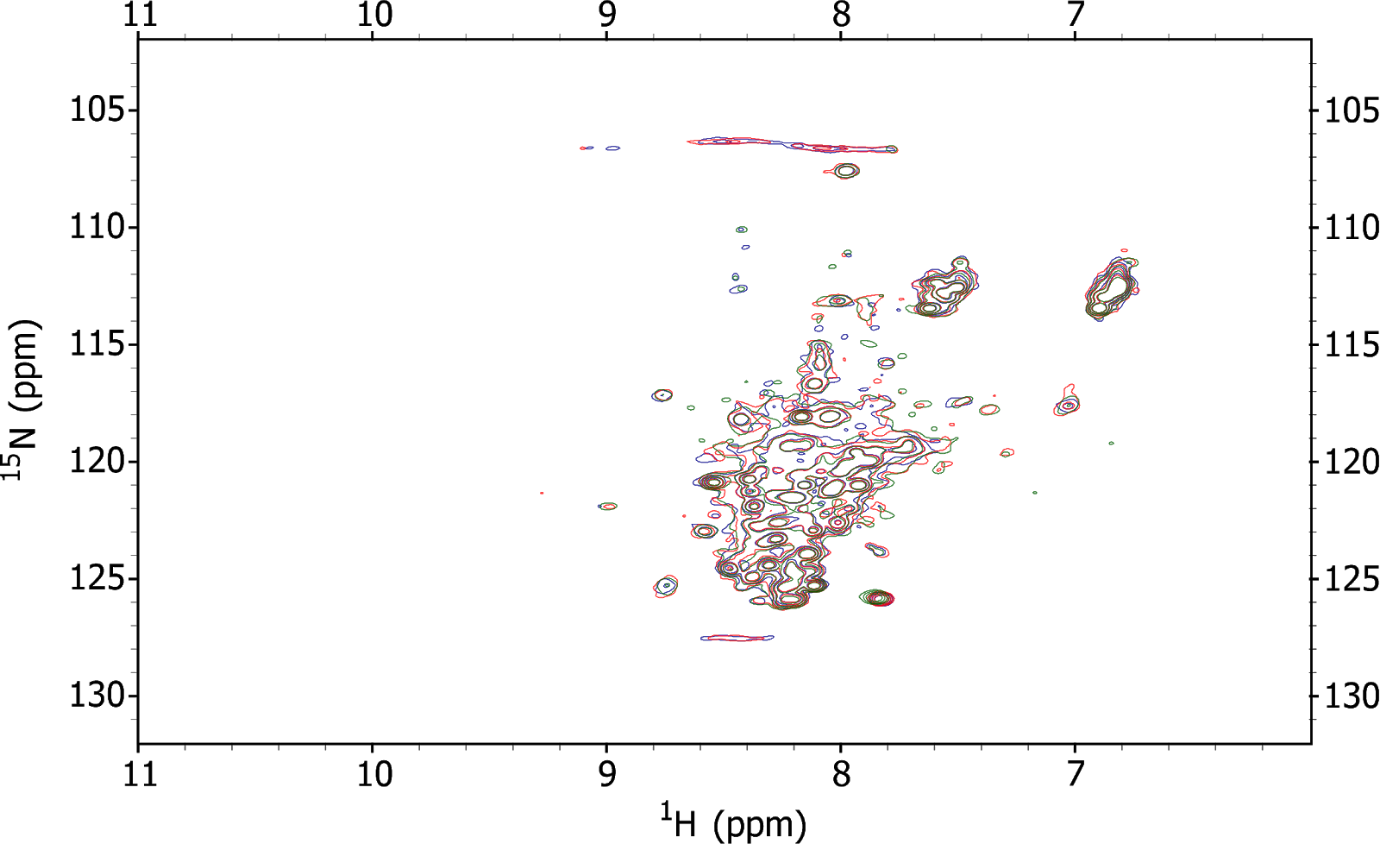

**Fig. S5:** Structural characterization of the ATP binding to DPCK variants by 2D NMR for

(A) DPCK-WT (30 μM), (B) DPCK-LH (100 μM) and (C) DPCK-M (100 μM) variants. Blue - protein, red - protein + ATP (1:1 molar ratio), green - protein + ATP (1:10 molar ratio). 2D NMR spectra were measured in 50 mM phosphate (pH 7.6), 280 mM NaCl, 20 mM KCl, 10 mM MgCl_2_, and 0.5 mM TCEP.

**Supplementary Table 1.** Selection of the target DPCK

| PDB code | 1UF9 | 2GRJ | **2IF2** |
| --- | --- | --- | --- |
| Organism | *Thermus thermophilus* | *Thermotoga*  *maritima* | *Aquifex*  *aeolicus* |
| Molecular weight, kDa | 22,64 | 21,79 | 24,0 |
| Length of sequence | 203 | 192 | 204 |
| Number of aromatic amino acids | 8  F - 4, Y - 1, W - 3 | 5  F - 3, Y - 1, W - 1 | 19  F - 6, Y - 12, W - 1 |
| Expression in *E. Coli* | High | High | High |
| Solubility | High | High | High |
| Oligomeric state | Monomeric | Oligomeric | Monomeric |
| Phosphotransferase activity | Yes | Yes | Yes |

**Supplementary Table 2**. Selection of DPCK variants lacking aromatic amino acids for further characterization (variants in red were excluded based on the solubility/activity profile)

| Protein | Expression in *E. Coli* | Solubility | Enzyme activity | |
| --- | --- | --- | --- | --- |
|  |  |  | ATPase | Phosphotransferase |
| DPCK-WT | High | High | No | Yes |
| DPCK-L | Low | Low | No | No |
| DPCK-LH | High | Low | Yes | Yes |
| DPCK-M | High | Medium | Yes | No |
| DPCK-MH | High | Medium | No | No |

**Supplementary Table 3**. The secondary structure content of the proteins

| Method | Deconvolutional analysis  of CD spectra | | | | Analysis of X-ray diffraction data  (PDB code: 2IF2) | | | | Analysis of NOESY spectra |
| --- | --- | --- | --- | --- | --- | --- | --- | --- | --- |
| Element | α-helix | β-sheet | β-turn | coil | α-helix | β-sheet | β-turn | coil | α-helix |
| DPCK-WT | 48 % | 9 % | 16 % | 28 % | 48 % | 12 % | 12 % | 14 % | 64 %* |
| DPCK-LH | 39 % | 11 % | 20 % | 30 % | — | — | — | — | 28 %* |
| DPCK-M | 44 % | 6 % | 21 % | 30 % | — | — | — | — | 7 %* |

* The content of α-helix was estimated by dividing the number of α-helical peaks estimated from NOESY spectra by the total number of amino acids in proteins.

***Gene sequences***

Wild Type (DPCK-WT)

ATGAAACGTATCGGTCTGACCGGTAACATCGGTTGCGGTAAATCTACCGTTGCGCAGATGTTCCGTGAACTGGGTGCGTACGTTCTGGACGCGGACAAACTGATCCACTCTTTCTACCGTAAAGGTCACCCGGTTTACGAAGAAGTTGTTAAAACCTTCGGTAAAGGTATCCTGGACGAAGAAGGTAACATCGACCGTAAAAAACTGGCGGACATCGTTTTCAAAGACGAAGAAAAACTGCGTAAACTGGAAGAAATCACCCACCGTGCGCTGTACAAAGAAATCGAAAAAATCACCAAAAACCTGTCTGAAGACACCCTGTTCATCCTGGAAGCGTCTCTGCTGGTTGAAAAAGGTACCTACAAAAACTACGACAAACTGATCGTTGTTTACGCGCCGTACGAAGTTTGCAAAGAACGTGCGATCAAACGTGGTATGTCTGAAGAAGACTTCGAACGTCGTTGGAAAAAACAGATGCCGATCGAAGAAAAAGTTAAATACGCGGACTACGTTATCGACAACTCTGGTTCTATCGAAGAAACCTACAAACAGGTTAAAAAAGTTTACGAAGAACTGACCCGTGACCCGCTGGAA

Leucine - His Mutant (DPCK-L)

ATGAAACGTATCGGTCTGACCGGTAACATCGGTTGCGGTAAATCTACCGTTGCGCAGATGCTGCGTGAACTGGGTGCGCTGGTTCTGGACGCGGACAAACTGATCCTGTCTCTGCTGCGTAAAGGTCTGCCGGTTCTGGAAGAAGTTGTTAAAACCCTGGGTAAAGGTATCCTGGACGAAGAAGGTAACATCGACCGTAAAAAACTGGCGGACATCGTTCTGAAAGACGAAGAAAAACTGCGTAAACTGGAAGAAATCACCCTGCGTGCGCTGCTGAAAGAAATCGAAAAAATCACCAAAAACCTGTCTGAAGACACCCTGCTGATCCTGGAAGCGTCTCTGCTGGTTGAAAAAGGTACCCTGAAAAACCTGGACAAACTGATCGTTGTTCTGGCGCCGCTGGAAGTTTGCAAAGAACGTGCGATCAAACGTGGTATGTCTGAAGAAGACCTGGAACGTCGTCTGAAAAAACAGATGCCGATCGAAGAAAAAGTTAAACTGGCGGACCTGGTTATCGACAACTCTGGTTCTATCGAAGAAACCCTGAAACAGGTTAAAAAAGTTCTGGAAGAACTGACCCGTGACCCGCTGGAA

Leucine + His Mutant (DPCK-LH)

ATGAAACGTATCGGTCTGACCGGTAACATCGGTTGCGGTAAATCTACCGTTGCGCAGATGCTGCGTGAACTGGGTGCGCTGGTTCTGGACGCGGACAAACTGATCCACTCTCTGCTGCGTAAAGGTCACCCGGTTCTGGAAGAAGTTGTTAAAACCCTGGGTAAAGGTATCCTGGACGAAGAAGGTAACATCGACCGTAAAAAACTGGCGGACATCGTTCTGAAAGACGAAGAAAAACTGCGTAAACTGGAAGAAATCACCCACCGTGCGCTGCTGAAAGAAATCGAAAAAATCACCAAAAACCTGTCTGAAGACACCCTGCTGATCCTGGAAGCGTCTCTGCTGGTTGAAAAAGGTACCCTGAAAAACCTGGACAAACTGATCGTTGTTCTGGCGCCGCTGGAAGTTTGCAAAGAACGTGCGATCAAACGTGGTATGTCTGAAGAAGACCTGGAACGTCGTCTGAAAAAACAGATGCCGATCGAAGAAAAAGTTAAACTGGCGGACCTGGTTATCGACAACTCTGGTTCTATCGAAGAAACCCTGAAACAGGTTAAAAAAGTTCTGGAAGAACTGACCCGTGACCCGCTGGAA

HotSpotWizard - His Mutant (DPCK-M)

ATGAAACGTATCGGTCTGACCGGTAACATCGGTTGCGGTAAATCTACCGTTGCGCAGATGCTGCGTGAACTGGGTGCGCCGGTTCTGGACGCGGACAAACTGATCCGTTCTGTTGTTCGTAAAGGTGGTCCGGTTCTGGAAGAAGTTGTTAAAACCGTTGGTAAAGGTATCCTGGACGAAGAAGGTAACATCGACCGTAAAAAACTGGCGGACATCGTTGTTAAAGACGAAGAAAAACTGCGTAAACTGGAAGAAATCACCTCTCGTGCGCTGCGTAAAGAAATCGAAAAAATCACCAAAAACCTGTCTGAAGACACCCTGGTTATCCTGGAAGCGTCTCTGCTGGTTGAAAAAGGTACCGACAAAAACGTTGACAAACTGATCGTTGTTGACGCGCCGGAAGAAGTTTGCAAAGAACGTGCGATCAAACGTGGTATGTCTGAAGAAGACGCGGAACGTCGTATCAAAAAACAGATGCCGATCGAAGAAAAAGTTAAACTGGCGGACGTTGTTATCGACAACTCTGGTTCTATCGAAGAAACCCGTAAACAGGTTAAAAAAGTTCTGGAAGAACTGACCCGTGACCCGCTGGAA

HotSpotWizard + His Mutant (DPCK-MH)

ATGAAACGTATCGGTCTGACCGGTAACATCGGTTGCGGTAAATCTACCGTTGCGCAGATGCTGCGTGAACTGGGTGCGCCGGTTCTGGACGCGGACAAACTGATCCACTCTGTTGTTCGTAAAGGTCACCCGGTTCTGGAAGAAGTTGTTAAAACCGTTGGTAAAGGTATCCTGGACGAAGAAGGTAACATCGACCGTAAAAAACTGGCGGACATCGTTGTTAAAGACGAAGAAAAACTGCGTAAACTGGAAGAAATCACCCACCGTGCGCTGCGTAAAGAAATCGAAAAAATCACCAAAAACCTGTCTGAAGACACCCTGGTTATCCTGGAAGCGTCTCTGCTGGTTGAAAAAGGTACCGACAAAAACGTTGACAAACTGATCGTTGTTGACGCGCCGGAAGAAGTTTGCAAAGAACGTGCGATCAAACGTGGTATGTCTGAAGAAGACGCGGAACGTCGTATCAAAAAACAGATGCCGATCGAAGAAAAAGTTAAACTGGCGGACGTTGTTATCGACAACTCTGGTTCTATCGAAGAAACCCGTAAACAGGTTAAAAAAGTTCTGGAAGAACTGACCCGTGACCCGCTGGAA

***Protein sequences***

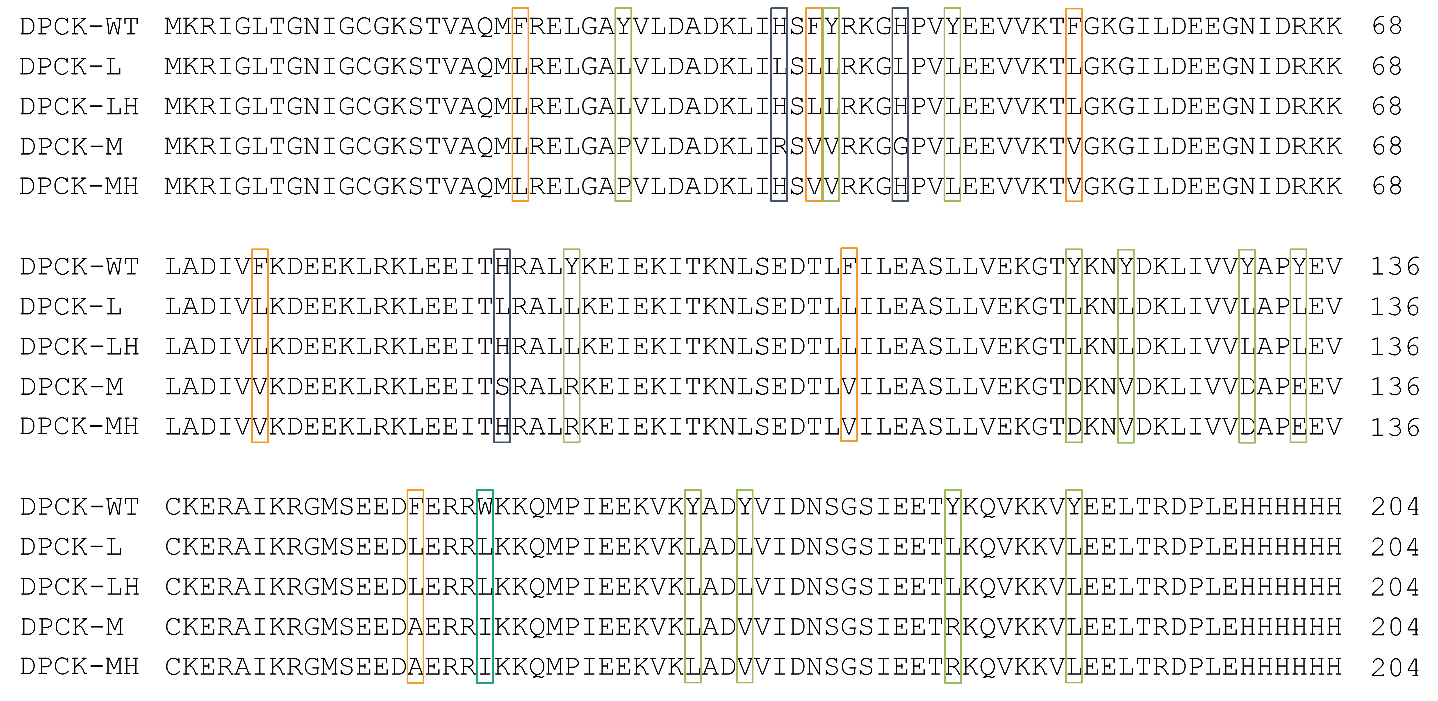
